# Epigenetic activation of CHST2 by promoter hypomethylation promotes progression of triple-negative breast cancer

**DOI:** 10.64898/2026.02.12.705678

**Authors:** Shukun Qu, Kangning Li, Lili Fan, Chenxue Miao, Weiyu Wang, Xu Wang, Guang Song

## Abstract

Epigenetic dysregulation is a hallmark of triple-negative breast cancer (TNBC), yet the oncogenic relevance of specific epigenetically activated genes remains poorly defined. In this study, we demonstrate that carbohydrate sulfotransferase 2 (CHST2) is aberrantly upregulated in TNBC across multiple transcriptomic analyses as a consequence of promoter hypomethylation. Elevated CHST2 expression is associated with aggressive clinicopathological characteristics and has been linked to unfavorable patient outcomes in independent cohorts. Functional analyses reveal that CHST2 primarily promotes TNBC cell migration and invasion, while genetic silencing of CHST2 markedly attenuates these malignant phenotypes. Importantly, mutational disruption of CHST2 catalytic activity abolishes its pro-migratory effects, indicating that sulfotransferase activity is essential for CHST2-driven invasiveness. Mechanistically, promoter hypomethylation is experimentally supported as a driver of CHST2 transcriptional activation, linking epigenetic deregulation to enhanced tumor aggressiveness. In addition, tumors with high CHST2 expression exhibit distinct immune-related transcriptional features within the tumor microenvironment. Collectively, these findings identify CHST2 as an epigenetically activated driver of invasive behavior in TNBC and highlight its potential value as a biomarker and therapeutic target for aggressive disease.

## Introduction

Triple-negative breast cancer (TNBC) is an aggressive and biologically heterogeneous subtype of breast cancer, clinically defined by the absence of estrogen receptor, progesterone receptor, and human epidermal growth factor receptor 2 expression(1, 2). Owing to its pronounced invasive potential, early metastatic dissemination, and lack of effective targeted therapies, TNBC continues to be associated with unfavorable clinical outcomes and poses a substantial therapeutic challenge(3).

Beyond genetic alterations, accumulating evidence highlights epigenetic dysregulation—particularly aberrant DNA methylation—as a critical determinant of transcriptional reprogramming and malignant behavior in TNBC(4). While widespread DNA methylation changes have been documented, it remains unclear which methylation-dependent events drive the activation of functionally relevant genes and how such epigenetic alterations contribute to tumor aggressiveness. Elucidating these mechanisms is essential for identifying epigenetically driven vulnerabilities in TNBC(5).

Among epigenetic mechanisms, DNA methylation at gene promoter regions constitutes an important layer contributing to transcriptional regulation(6). In cancer, promoter hypomethylation is frequently associated with aberrant gene activation, enabling tumor cells to acquire enhanced migratory and invasive capacities and to adapt to the tumor microenvironment(7, 8). Although extensive efforts have documented methylation-associated transcriptional changes in breast cancer, accumulating evidence suggests that only a fraction of these alterations function as bona fide epigenetic drivers that actively shape malignant phenotypes(5). Identifying such functionally relevant events, and distinguishing them from secondary epigenetic changes, remains a critical challenge in cancer biology, particularly in aggressive breast cancer subtypes such as TNBC.

Glycosylation represents a major post-translational modification influencing protein stability, receptor signaling, and cell–cell interactions, thereby shaping tumor–microenvironment communication(9). Aberrant regulation of glycosylation-associated enzymes has been increasingly implicated in cancer progression and metastatic dissemination(10). Among these enzymes, the carbohydrate sulfotransferase (CHST) family catalyzes the sulfation of glycosaminoglycans and glycoproteins, modulating extracellular matrix organization and signaling pathways associated with tumor aggressiveness(11). Although several CHST family members have been linked to malignant phenotypes in diverse cancer types, the biological role of CHST2 in triple-negative breast cancer remains poorly defined(12). Moreover, whether CHST2 expression is subject to epigenetic regulation and functionally contributes to TNBC progression has not been systematically investigated.

In this study, we sought to identify DNA methylation–driven regulators with functional relevance in triple-negative breast cancer through integrated analyses of transcriptomic and DNA methylation profiles. Focusing on carbohydrate sulfotransferase 2 (CHST2), we investigated its expression pattern, epigenetic regulation, and contribution to malignant phenotypes in TNBC. By combining epigenomic analyses with functional and mechanistic approaches, this work aims to elucidate how aberrant epigenetic activation of glycan-modifying enzymes influences tumor aggressiveness and to provide insight into the biological significance of epigenetically driven metabolic reprogramming in breast cancer.

## Materials and Methods

### Data collection and preprocessing

Transcriptomic data (HTSeq-TPM) and the corresponding clinicopathological information of breast cancer patients were obtained from The Cancer Genome Atlas (TCGA) database(13). DNA methylation data generated using the Illumina HumanMethylation450 BeadChip platform were downloaded for the same cohort. Triple-negative breast cancer (TNBC) samples were defined based on negative expression of estrogen receptor (ER), progesterone receptor (PR), and HER2.

To validate the robustness of gene expression patterns, an independent breast cancer cohort (GSE96058) was retrieved from the Gene Expression Omnibus (GEO) database(14). The expression and promoter methylation status of CHST2 in breast cancer cell lines were analyzed using data from the Cancer Cell Line Encyclopedia (CCLE)(15). Genes with zero counts in more than 50% of samples were removed to reduce technical noise. H3K27ac ChIP–seq datasets of breast cancer cell lines were obtained from GEO (GSE85158 and GSE69107)(16, 17).

### Differential gene expression analysis

Differential expression analysis between TNBC and non-TNBC samples was performed using the R package DESeq2(v1.50.2) based on raw count matrices. Genes with an adjusted p value (False discovery rate, FDR) < 0.05 and |log₂ fold change| > 1 were considered differentially expressed. Multiple testing correction was conducted using the Benjamini–Hochberg method. Visualization of differentially expressed genes (DEGs) was performed using the ggplot2 package.

### DNA methylation analysis

TCGA breast cancer DNA methylation data were downloaded using the TCGAbiolinks package(v2.38.0). Probes and samples with more than 30% missing values were excluded. Remaining missing values were imputed using the *impute.knn* function from the impute package(). Promoter regions were defined as ±1 kb surrounding the transcription start site (TSS). The average β value of CpG probes located within promoter regions was used to represent promoter methylation levels. Differential promoter methylation analysis between TNBC and non-TNBC samples was performed using the limma package. Promoters with |Δβ| > 0.1–0.2 and FDR < 0.05 were considered significantly differentially methylated.

Reduced representation bisulfite sequencing (RRBS) data of breast cancer cell lines were downloaded from the Sequence Read Archive (SRA). Raw data were processed into methylation matrices using BS-Seeker2(v2.1.8) according to the official pipeline. Downstream filtering, imputation, and differential analysis were conducted using the same criteria as described above.

### Integration of gene expression and DNA methylation

Genes with significantly altered promoter methylation were defined as genes harboring differentially methylated promoters (DMPs). These genes were subsequently intersected with DEGs to generate four gene categories: upregulated–hypermethylated, upregulated–hypomethylated, downregulated–hypermethylated, and downregulated–hypomethylated. Intersections were visualized using Venn diagrams and volcano plots.

Spearman correlation analysis was performed to assess the association between promoter methylation levels and gene expression. Genes with |r| > 0.3 and FDR < 0.05 were considered significantly correlated.

### Functional enrichment and gene set enrichment analysis

Gene Ontology (GO) enrichment analysis was performed using the clusterProfiler package(v4.18.4) (enrichGO function) for biological process (BP) terms. Gene sets with an adjusted p < 0.05 were considered significantly enriched (minGSSize = 20, maxGSSize = 500). Enriched terms were ranked by −log₁₀(*p* value). For gene set enrichment analysis (GSEA), gene symbols were converted to Entrez IDs using the bitr function. Genes were ranked according to log₂ fold change and analyzed using the gseGO function. Gene sets with FDR < 0.05 were considered significantly enriched (minGSSize = 10, maxGSSize = 500).

### Immune infiltration and ESTIMATE analysis

Single-sample gene set enrichment analysis (ssGSEA) was performed using the GSVA package (v2.4.4,gsva function; method = “ssgsea”, kcdf = “Gaussian”) to estimate immune cell infiltration scores for each sample. Samples were divided into high- and low-CHST2 expression groups based on the median expression level. Differences in immune cell infiltration between groups were visualized using boxplots. Spearman correlation analysis was performed to evaluate the associations between CHST2 expression and immune cell infiltration scores, and the results were visualized using lollipop charts. Tumor stromal and immune components were evaluated using the ESTIMATE package(v1.0.13). Correlations between CHST2 expression and StromalScore, ImmuneScore, and ESTIMATEScore were analyzed and visualized using the ggpubr package.

### Single-cell RNA sequencing analysis

Single-cell RNA sequencing data (GSE176078) were downloaded from GEO database(18) and analyzed using the Seurat package(v5.4.0). Seurat objects were constructed using the CreateSeuratObject function. Quality control was performed to retain high-quality cells with 300–8000 detected genes, total UMI counts < 100,000, mitochondrial gene proportion < 20%, and hemoglobin gene proportion < 5%. Potential doublets were identified and removed using the scDblFinder package(v1.23.2). After quality control and doublet removal, data were normalized using the NormalizeData function, and 3000 highly variable genes were identified using FindVariableFeatures. To correct for batch effects, data integration was performed using the FindIntegrationAnchors and IntegrateData functions. Dimensionality reduction was conducted using principal component analysis (PCA), followed by UMAP visualization. Cell clustering was performed using the FindNeighbors and FindClusters functions, and clusters were visualized using DimPlot. Gene set scoring at the single-cell level was conducted using the AddModuleScore function, and the results were visualized using violin plots.

### H3K27ac ChIP–seq data processing

Raw H3K27ac ChIP–seq data were downloaded from the SRA database and converted to FASTQ format using parallel-fastq-dump(v0.6.7). Quality control was performed using FastQC, and adapter trimming and quality filtering were conducted using fastp(v2.5.4). Clean reads were aligned to the human reference genome (hg38) using Bowtie2(v2.5.4). PCR duplicates were marked and removed using Picard and SAMtools(v1.17). Peak calling was performed using MACS2(v3.0.3), with H3K27ac ChIP samples as treatment and corresponding input samples as controls. Signal tracks were converted to BigWig format for visualization using bedGraphToBigWig(v482).

### Cell culture and transfection

Human breast cancer cell lines MDA-MB-231 and MCF7 were obtained from Procell Life Science & Technology (Wuhan, China) and cultured in Dulbecco’s Modified Eagle Medium (DMEM) with 10% fetal bovine serum (FBS) and 1% penicillin–streptomycin at 37°C with 5% CO₂. Human CHST2 cDNA was cloned into the lentiviral vector pCDH-CMV-MCS-EF1-Puro-3×Flag. A catalytically inactive mutant (N475A) was generated using PCR-based site-directed mutagenesis. Lentiviral shRNA targeting CHST2 was purchased from YouBio Biotech (Changsha, China). Stable cell lines were selected using puromycin (1 μg/mL).

### Cell proliferation assays

Cell proliferation was assessed using the Cell Counting Kit-8 (CCK-8) assay. Cells were seeded into 96-well plates at a density of 1 × 10⁴ cells per well, with five technical replicates per group, and cultured at 37°C in a humidified incubator containing 5% CO₂. At 0, 24, 48, and 72 h, the culture medium was removed, and cells were gently washed twice with phosphate-buffered saline. Subsequently, 100 μL of DMEM containing 10% CCK-8 reagent was added to each well and incubated for 1 h. Absorbance was measured at 450 nm using a microplate reader. Relative cell proliferation was calculated after subtraction of background absorbance.

### Transwell migration and invasion assays

Cell migration and invasion were evaluated using Transwell chambers (8-μm pore size). Cells were trypsinized, resuspended in serum-free DMEM, and adjusted to a concentration of 1 × 10⁵ cells/mL. For migration assays, 200 μL of cell suspension was added to the upper chamber, while 800 μL of complete medium containing 20% fetal bovine serum was placed in the lower chamber as a chemoattractant. After incubation at 37 °C for 24 h, non-migrated cells on the upper surface of the membrane were gently removed. Migrated cells were fixed with 4% paraformaldehyde for 30 min and stained with crystal violet for 15 min. For invasion assays, the upper chambers were pre-coated with Matrigel diluted 1:50 in serum-free DMEM and incubated at 37 °C for 2 h to allow gel formation. The subsequent procedures were identical to those described for migration assays. Representative images were captured under a light microscope, and cells were quantified using ImageJ software.

### Wound-healing assay

Wound-healing assays were performed to assess cell migratory capacity. Cells were seeded into six-well plates at a density of 5 × 10⁵ cells per well and cultured until reaching complete confluence. Linear wounds were generated using a 200-μL sterile pipette tip along predefined reference lines to ensure consistent imaging fields. Detached cells were removed by washing with phosphate-buffered saline, and cells were cultured in DMEM supplemented with 1% fetal bovine serum. Images were acquired at 0 and 24 h under identical microscopic fields. Wound closure was quantified using ImageJ software based on the change in wound area.

### RNA extraction and quantitative real-Time PCR

Total RNA was extracted using TRIzol reagent, following the manufacturer’s protocol. and cDNA was synthesized using a reverse transcription kit. Quantitative real-time PCR (RT–qPCR) was performed using SYBR Green chemistry. Relative expression levels were calculated using the 2⁻^ΔΔCt^ method with β-actin as the internal control.

### Western blotting

Total cellular proteins were extracted using lysis buffer (25 mM Tris–HCl (pH 7.6), 150 mM NaCl, 1% NP-40, 1% sodium deoxycholate, and 0.1% SDS) supplemented with phenylmethylsulfonyl fluoride (PMSF). Equal amounts of protein were separated by SDS–PAGE and transferred onto polyvinylidene fluoride (PVDF) membranes. Membranes were blocked with 5% non-fat milk and incubated with primary antibodies at 4 °C overnight, followed by incubation with horseradish peroxidase–conjugated secondary antibodies. All primary antibodies were purchased from Upingbio Biology (Hangzhou, China). Protein bands were visualized using enhanced chemiluminescence (ECL) reagents. Band intensities were quantified using ImageJ software and normalized to background signals and internal loading controls. The antibodies used for Western blot analysis were as follows: CHST2 antibody (YT0920, Immunoway, Plano, TX, USA), anti-Flag antibody (ab1162, Abcam, Cambridge, UK), GAPDH antibody (60004-1-lg, Proteintech, Chicago, IL, USA), β-actin antibody (81115-1-RR, Proteintech, Chicago, IL, USA), HRP-conjugated goat anti-rabbit IgG (SA00001-2, Proteintech, Chicago, IL, USA), and HRP-conjugated goat anti-mouse IgG (SA00001-1, Proteintech, Chicago, IL, USA).

### Bisulfite sequencing

Genomic DNA was extracted from MDA-MB-231 and MCF7 cells using a DNA extraction kit (DP304, Tiangen Biotech, Beijing, China). A total of 2 μg genomic DNA was subjected to bisulfite conversion using a bisulfite conversion kit (D0068M, Beyotime Biotechnology, Shanghai, China) according to the manufacturer’s instructions. BSP primers targeting the CHST2 promoter region were designed using MethPrimer, and primer sequences are listed in Table 1. Bisulfite-converted DNA was amplified by PCR using CHST2 promoter–specific primers. PCR products were purified and cloned into TA vectors using a TA cloning kit (ZC206, ZOMANBIO, Beijing, China). At least 6 individual clones per sample were randomly selected for Sanger sequencing (Sangon Biotech Co., Ltd, Shanghai, China). Sequencing results were analyzed using SnapGene (San Diego, California) to determine the methylation status at individual CpG sites.

### DNA demethylation treatment

To investigate the role of DNA methylation in regulating CHST2 expression, MCF7 cells were treated with 5-aza-2′-deoxycytidine (5 μmol/mL) for 24 h. Following treatment, genomic DNA was extracted and analyzed by BSP to assess changes in methylation levels within the CHST2 promoter region. Total RNA and protein were subsequently isolated for RT–qPCR and Western blot analyses to evaluate CHST2 expression at the transcriptional and protein levels.

### Statistical analysis

Statistical analyses were performed using R (v4.5.1) and GraphPad Prism (v9.0). All experiments were independently performed at least three times. Quantitative data are presented as mean ± standard deviation (SD). Statistical analyses for in vitro experiments were conducted using GraphPad Prism (v 9.0), unless otherwise specified. Comparisons between two groups were performed using Student’s *t*-test, while comparisons among multiple groups were analyzed by one-way analysis of variance (ANOVA) followed by appropriate post hoc tests.

Correlation analyses were conducted using Spearman’s rank correlation coefficient. For high-throughput data analyses, including transcriptomic and epigenomic datasets, multiple testing correction was applied using the false discovery rate (FDR) method where applicable. Densitometric analyses of Western blot and other image-based assays were performed using ImageJ software prior to statistical evaluation. A *p* value < 0.05 was considered statistically significant.

## Results

### Promoter hypomethylation contributes to transcriptional activation in TNBC

To systematically identify genes with aberrant expression in TNBC may be associated with promoter DNA methylation alterations, we integrated transcriptomic and DNA methylation data from TCGA breast cancer tissues (135 TNBC and 603 non-TNBC). Differential expression analysis identified 7,114 altered genes in TNBC, comprising 3,986 upregulated and 3,128 downregulated (Fig.1A). In parallel, promoter-level methylation analysis identified 1,880 genes with significant methylation changes, including 1,110 hypomethylated and 770 hypermethylated promoters in TNBC (Fig.1B).

**Fig. 1.**
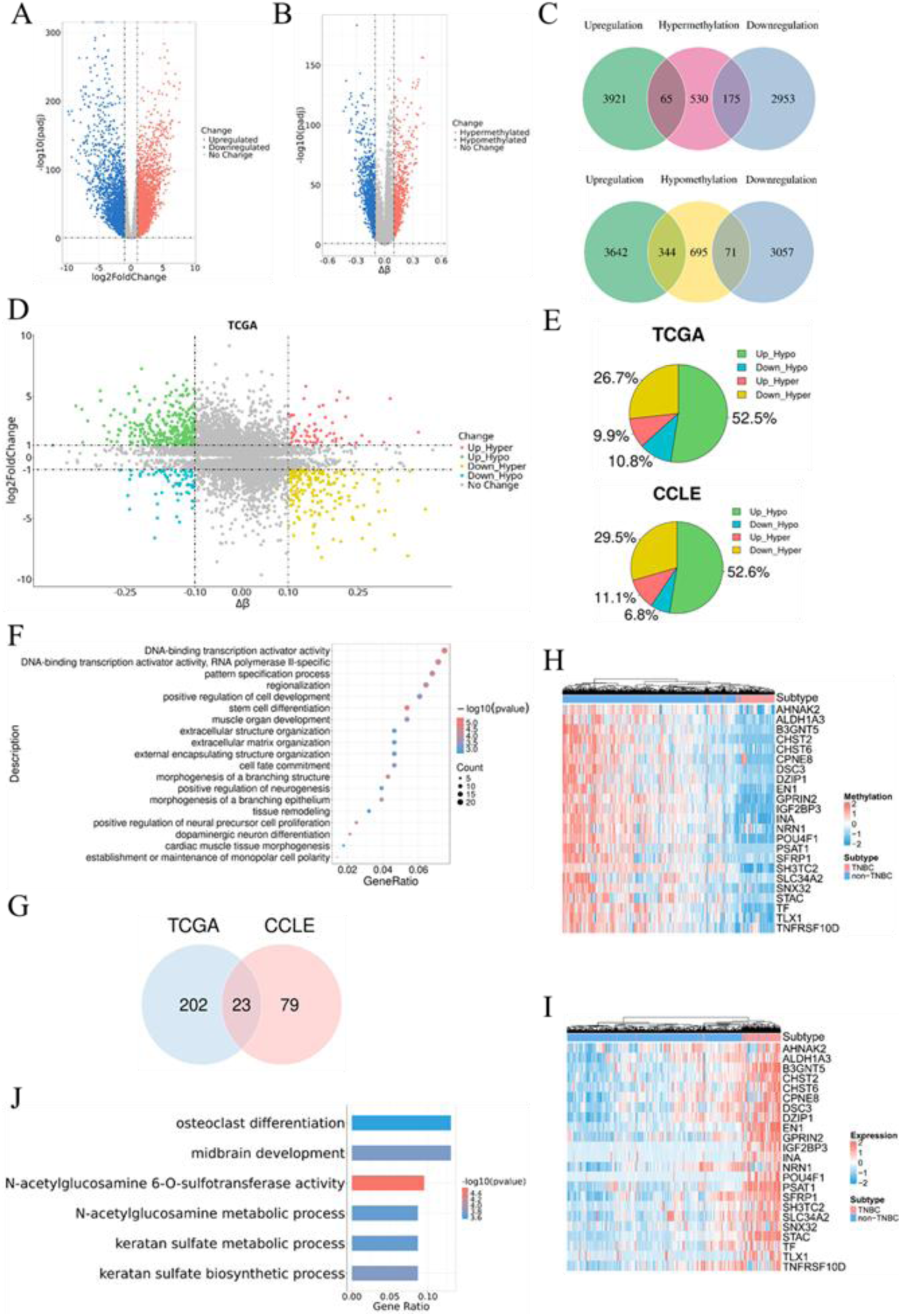
Promoter hypomethylation–associated gene activation in TNBC. (A) Volcano plot of differentially expressed genes (DEGs) between TNBC and non-TNBC samples in the TCGA cohort. Red and blue dots indicate significantly upregulated and downregulated genes in TNBC. (B) Volcano plot of gene-level promoter methylation differences between TNBC and non-TNBC samples in TCGA. Red dots denote hypermethylated promoters and blue dots denote hypomethylated promoters in TNBC. (C) Venn diagrams showing the overlap between genes with differential promoter methylation (DMGs) and genes that are upregulated or downregulated in TNBC in the TCGA cohort. (D) Integrated analysis of DEGs and DMGs classifying genes into five regulatory patterns: Up_Hyper, Up_Hypo, Down_Hyper, Down_Hypo, and No Change. (E) Proportions of the four major epigenetic regulatory patterns in TCGA and CCLE datasets. (F) Gene Ontology (GO) enrichment analysis of genes exhibiting the Up_Hypo pattern in TCGA. Bubble size indicates gene count and color reflects statistical significance. (G) Venn diagram showing the overlap of significantly negatively correlated Up_Hypo genes between TCGA and CCLE datasets. (H–I) Heatmaps showing expression levels (H) and promoter methylation levels (I) of 23 key Up_Hypo genes in TNBC versus non-TNBC samples from TCGA. (J) GO enrichment analysis of the 23 Up_Hypo key genes.

Intersection analysis between differentially expressed genes (DEGs) and differentially methylated promoters (DMPs) revealed distinct methylation–expression regulatory patterns (Fig.1C). Genes were classified into major categories: upregulated–hypermethylated (Up_Hyper), upregulated–hypomethylated (Up_Hypo), downregulated–hypermethylated (Down_Hyper), and downregulated–hypomethylated (Down_Hypo) (Fig.1D). Among these, the Up_Hypo group was the most prevalent (52.5%) (Fig.1E). This regulatory pattern was further examined using RNA-seq and RRBS data from the Cancer Cell Line Encyclopedia (CCLE), including 17 TNBC and 19 non-TNBC cell lines. Consistent with TCGA results, Up_Hypo genes remained the largest fraction (52.6%), supporting conservation of this epigenetic activation pattern across tumor tissues and cell line models (Fig. 1E and Fig. S1A–D).

Gene Ontology enrichment analysis of Up_Hypo genes from both TCGA and CCLE consistently showed enrichment in transcriptional activation, stemness maintenance, neurodevelopmental regulation, and extracellular matrix remodeling, implicating their roles in aggressive tumor phenotypes (Fig.1F and Fig.S1E). Further integration of Up_Hypo genes across TCGA and CCLE identified a core set of 23 genes that consistently exhibited promoter hypomethylation accompanied by elevated expression (Fig.1G). Unsupervised clustering based on these genes could robustly distinguished TNBC from non-TNBC, defining a TNBC-specific epigenetic landscape (Fig.1H-I and Fig.S1F-G). Further functional enrichment analysis of these 23 genes revealed that N-acetylglucosamine 6-O-sulfotransferase activity was the significantly enriched, with lowest adjusted *p* value (Fig.1J), implicating dysregulated carbohydrate sulfation pathways in TNBC. This observation prompted us to further interrogate CHST2 and CHST6, two epigenetically activated CHST family members within this core set.

### Epigenetic profiling of CHSTs identifies CHST2 as a key candidate in TNBC

Analysis of TCGA samples showed that CHST2 and CHST6 were both significantly upregulated in TNBC compared with non-TNBC tumors and exhibited reduced promoter methylation (Fig. 2A–B and Fig. 2D–E). For both genes, promoter methylation levels were inversely correlated with mRNA expression; however, this association was substantially stronger for CHST2 than for CHST6 (Fig. 2C and Fig. 2F). Similar expression and methylation patterns were observed in CCLE datasets (Fig. S2A–F).

**Fig. 2.**
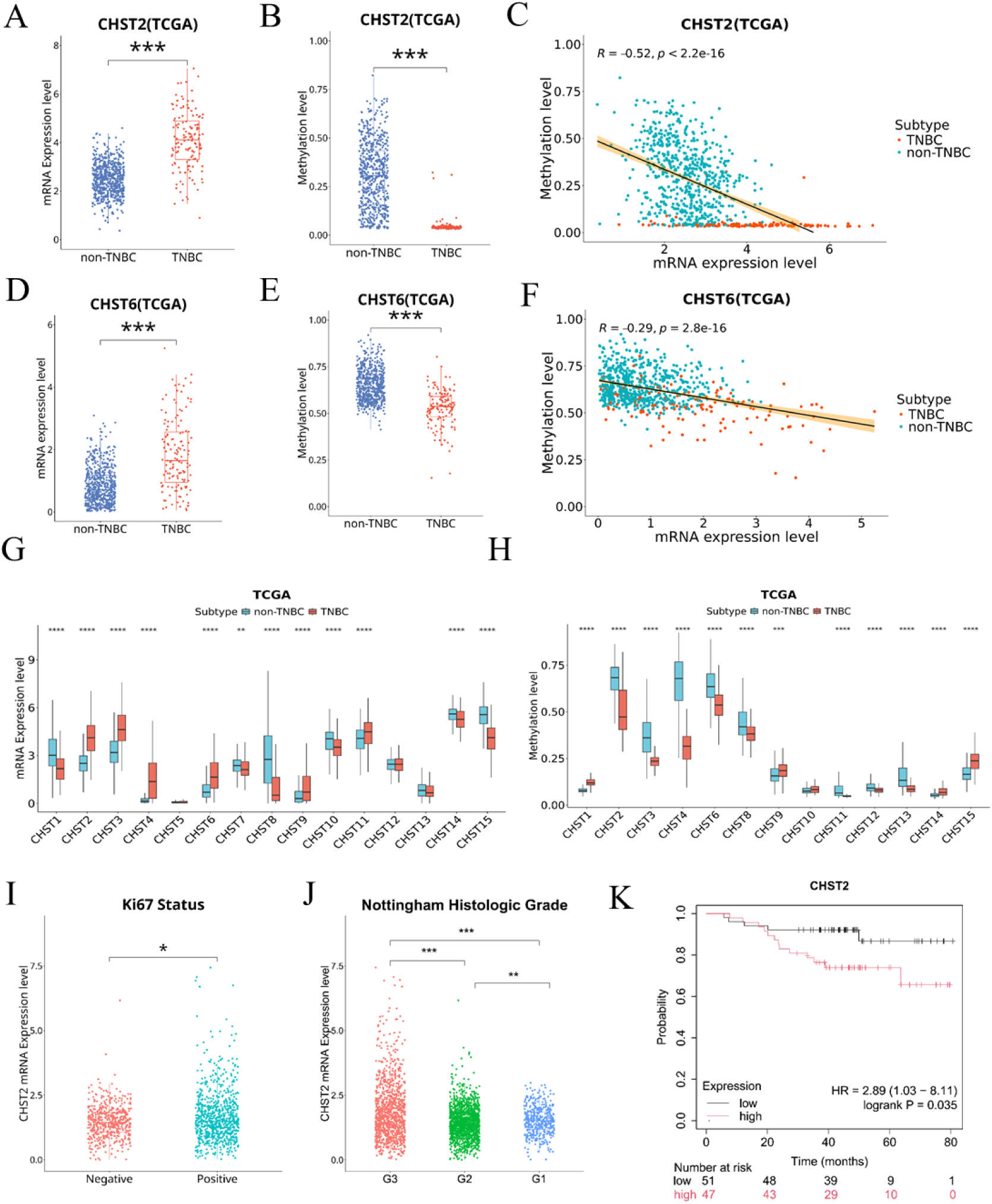
Epigenetic and clinical characterization of CHST family members in TNBC. (A–B) Box plots showing CHST2 mRNA expression (A) and its promoter methylation (B) in TNBC and non-TNBC samples from TCGA. (C) Correlation analysis between CHST2 mRNA expression and its promoter methylation in TCGA samples. Each dot represents one tumor; TNBC and non-TNBC samples are indicated by different colors. (D–E) Box plots showing CHST6 mRNA expression (D) and its promoter methylation (E) in TNBC and non-TNBC samples from TCGA. (F) Correlation analysis between CHST6 mRNA expression and its promoter methylation in TCGA samples. (G) Box plots summarizing differential expression of CHST family members in TNBC versus non-TNBC samples from TCGA. (H) Box plots showing promoter methylation differences of CHST family genes between TNBC and non-TNBC samples in TCGA. (I) CHST2 expression levels in Ki-67–positive and Ki-67–negative breast cancer samples from the GSE96058 cohort. (J) CHST2 expression levels across different Nottingham histologic grades in the GSE96058 cohort. (K) Kaplan–Meier survival analysis showing the association between CHST2 expression and overall survival in TNBC patients based on the Kaplan–Meier Plotter database. * *p* <0.05; ** *p* <0.01; *** *p* <0.001.

We next performed a systematic analysis of the CHST gene family. In TCGA tissue samples CHST2, CHST3, CHST4, and CHST6 were highly expressed in TNBC and were accompanied by promoter hypomethylation (Fig. 2G–H). For CCLE cell lines, However, only CHST2 and CHST6 retained this concordant Up_Hypo pattern (Fig. S2G–H), indicating that the methylation–expression coupling observed in primary tumors is not uniformly preserved across all CHST members in vitro. Moreover, Analysis of an independent cohort (GSE96058) confirmed elevated expression of CHST2, CHST3 and CHST4 in TNBC (Fig. S2I-J), supporting transcriptional reproducibility across datasets. Collectively, CHST2 represents the most consistently and robustly Up_Hypo-regulated member of the CHST family across both tumor tissues and breast cancer cell line models.

To assess the clinical relevance of CHST2, we analyzed the GSE96058 cohort. CHST2 expression was higher in Ki-67–positive tumors compared with Ki-67–negative tumors (Fig. 2I) and increased with higher Nottingham histologic grade (Fig. 2J). Kaplan–Meier survival analysis further revealed that elevated CHST2 expression was associated with poorer overall survival in patients with TNBC (Fig. 2K). It should be noted that similar associations were not statistically significant in the TCGA cohort (data not shown). Collectively, these findings highlight CHST2 as the most prominently epigenetically activated CHST gene in TNBC with potential clinical relevance.

### Single-cell analysis reveals epithelial-enriched CHST2 linked to EMT in TNBC

To delineate the cellular distribution of CHST2 across breast cancer subtypes, we analyzed single-cell RNA-seq data from primary tumors in the GSE176078 cohort. After quality control, doublet removal, and batch correction, cells were classified into seven major cell types based on canonical marker genes (Fig. 3A; Fig. S3A–H). CHST2 expression displayed a pronounced subtype-dependent pattern. In TNBC, CHST2 expression was enriched in epithelial cells, whereas in non-TNBC tumors it was preferentially detected in endothelial and mesenchymal compartments (Fig. 3B). Accordingly, the proportion of CHST2-positive epithelial cells was significantly higher in TNBC than in non-TNBC samples (Fig. 3C).

**Fig. 3.**
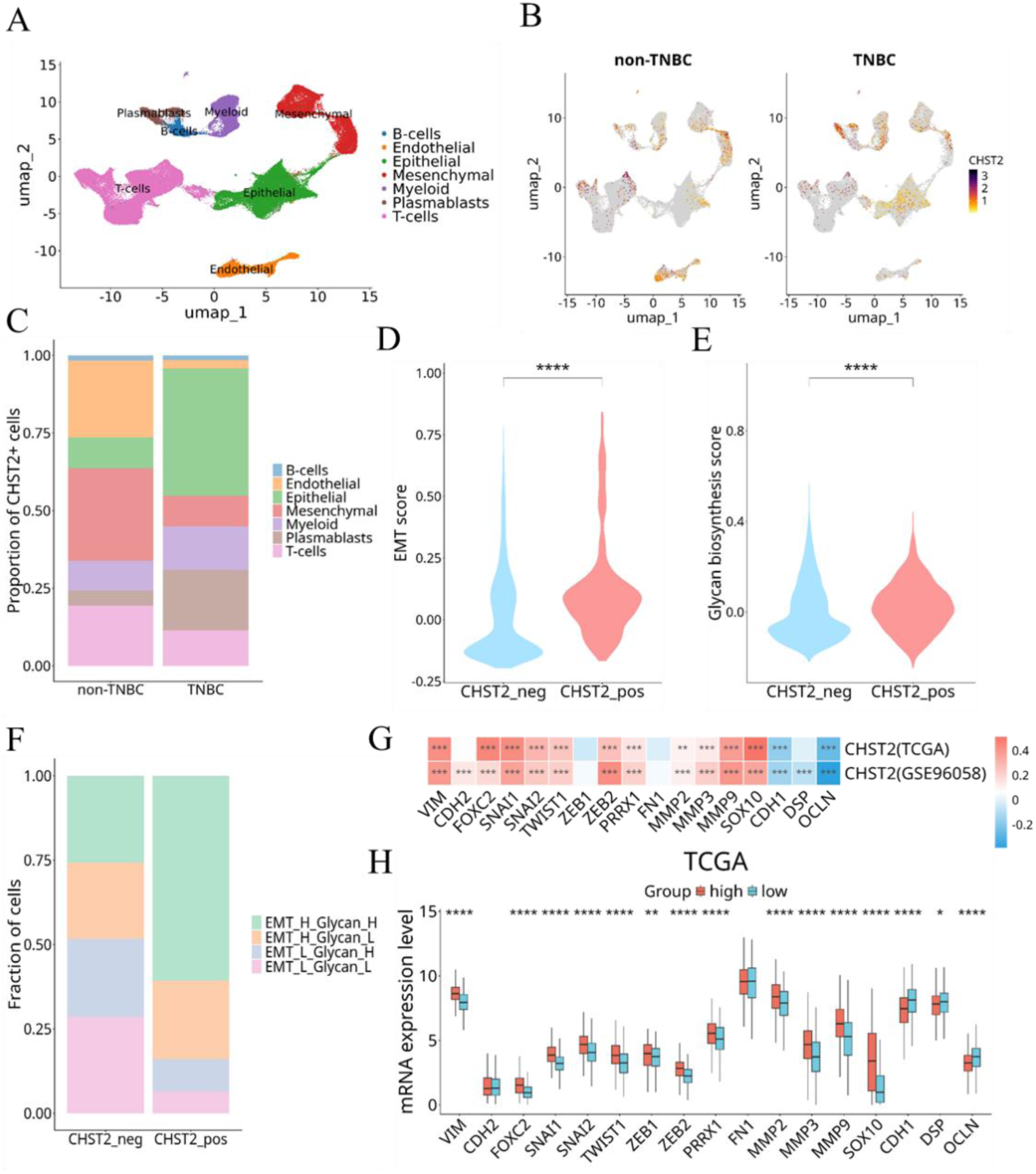
Single-cell analysis links CHST2 to EMT and glycan biosynthesis in TNBC. (A) UMAP visualization showing the distribution of major cell types in breast cancer samples based on single-cell RNA sequencing data. (B) UMAP feature plots illustrating CHST2 expression across different cell types in TNBC and non-TNBC samples. (C) Proportions of CHST2-positive cells among different cell types in TNBC and non-TNBC samples. (D) Comparison of epithelial–mesenchymal transition scores between CHST2-positive and CHST2-negative epithelial cells in TNBC samples. (E) Comparison of glycan biosynthesis scores between CHST2-positive and CHST2-negative epithelial cells in TNBC samples. (F) Distribution of epithelial cell subpopulations stratified by EMT and glycan biosynthesis scores in CHST2-positive and CHST2-negative cells from TNBC samples. Cells were classified as EMT_H or EMT_L, and Glycan_H or Glycan_L based on whether the corresponding scores were above or below the median. (G) Correlation analysis between CHST2 expression and EMT-related genes in the TCGA and GSE96058 datasets. (H) Comparison of EMT-related gene expression between CHST2-high and CHST2-low groups in the TCGA cohort. Samples were stratified according to the median CHST2 expression level. * *p* <0.05; ** *p* <0.01; *** *p* <0.001; **** *p* <0.0001.

To determine whether CHST2 expression marks aggressive epithelial states in TNBC, we scored epithelial cells using a canonical EMT gene signature comprising established mesenchymal markers (VIM, CDH2, FOXC2, SNAI1, SNAI2, TWIST1, ZEB1, ZEB2, PRRX1, FN1, MMP2, MMP3, MMP9, and SOX10) and epithelial markers (CDH1, DSP, and OCLN)(19, 20). CHST2-positive epithelial cells displayed significantly higher EMT scores than CHST2-negative cells (Fig. 3D).

Given that CHST2 catalyzes glycan sulfation, we next assessed whether CHST2-positive cells also exhibit enhanced upstream glycan biosynthetic activity. A glycan biosynthesis signature encompassing key enzymes involved in fucosylation(FUT6 and FUT7), sialylation(ST3GAL4 and ST3GAL6), and glycan elongation(B4GALT1 and B3GNT5) was applied to TNBC epithelial cells(21–24). CHST2-positive cells showed significantly elevated glycan biosynthesis scores compared with CHST2-negative cells (Fig. 3E). Stratification by EMT and glycan scores further demonstrated that CHST2-positive cells were enriched in EMT-high and glycan-high subpopulations (Fig. 3F).

At the bulk transcriptomic level, CHST2 expression positively correlated with mesenchymal markers and inversely correlated with epithelial markers in both TCGA and GSE96058 cohorts (Fig. 3G). Consistently, CHST2-high tumors exhibited increased mesenchymal marker expression and reduced epithelial marker expression across datasets (Fig. 3H; Fig. S3I).

Collectively, these findings link preferential CHST2 expression in TNBC epithelial cells to EMT activation and enhanced glycan biosynthetic programs, supporting a tumor cell–intrinsic association between CHST2 and aggressive TNBC phenotypes.

### CHST2 expression correlates with immune and interferon features in TNBC

To investigate the association between CHST2 expression and the tumor immune context, TCGA breast cancer samples were stratified into CHST2-high and CHST2-low groups based on median expression (Fig. 4A). Differential expression analysis identified 2,894 CHST2-associated genes, predominantly upregulated in CHST2-high tumors. Gene set enrichment analysis revealed significant enrichment of immune-related programs, including interferon signaling, cytokine–cytokine receptor interaction, and hematopoietic lineage pathways (Fig. 4B), indicating an immune-associated transcriptional phenotype.

**Fig. 4.**
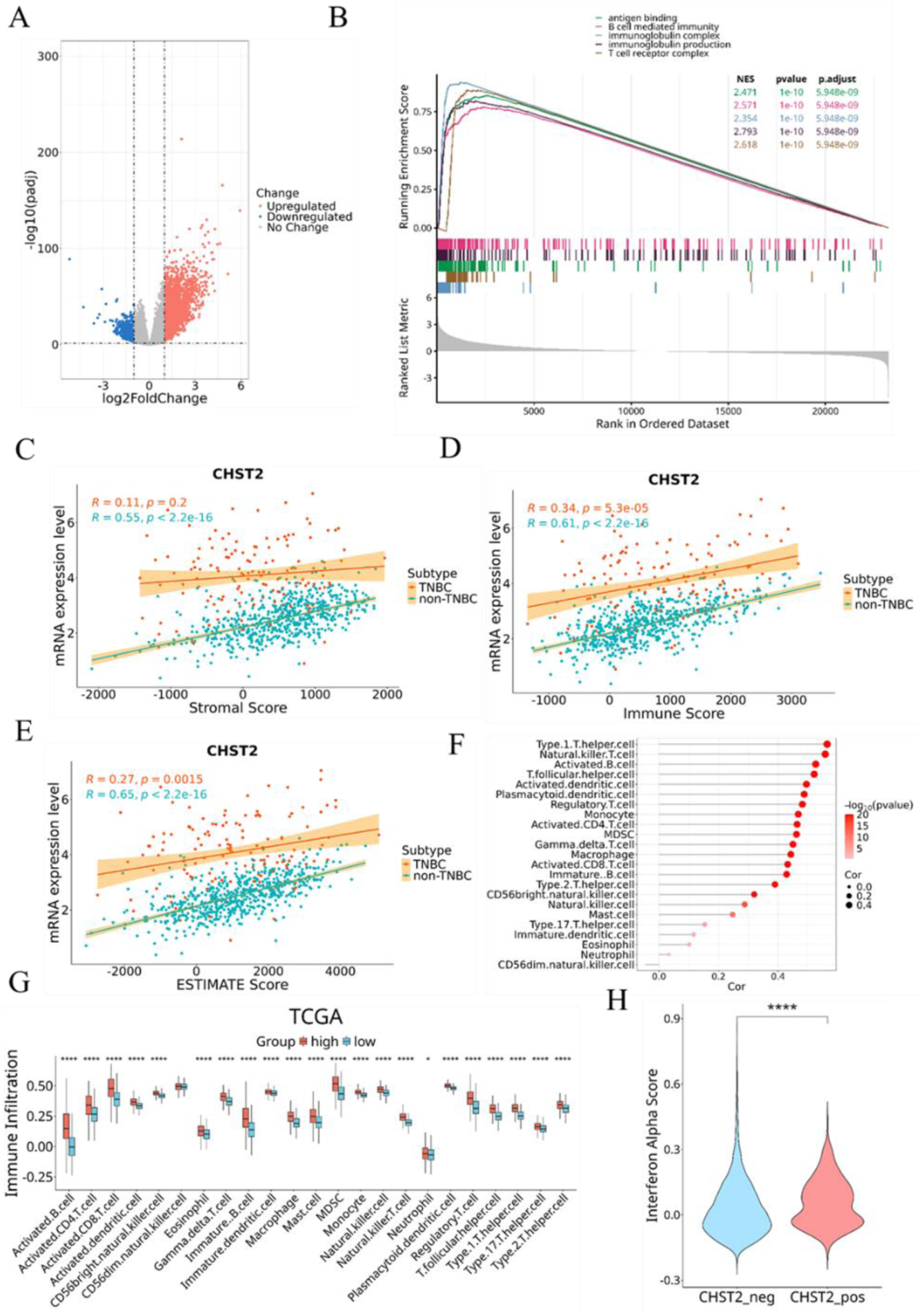
CHST2 expression is associated with stromal, immune, and interferon features in the TNBC microenvironment. (A) Differentially expressed genes between CHST2-high and CHST2-low groups in the TCGA cohort. Samples were stratified according to the median CHST2 expression level. (B) Gene set enrichment analysis of differentially expressed genes between CHST2-high and CHST2-low groups in TCGA, showing the top five pathways ranked by normalized enrichment score. (C–E) Correlation analysis between CHST2 expression and stromal score (C), immune score (D), and ESTIMATE score (E) in TCGA samples. Each dot represents one tumor sample; TNBC and non-TNBC samples are indicated by different colors. Correlation coefficients and *p* values were calculated using Spearman’s correlation analysis. (F) Lollipop plot showing correlations between CHST2 expression and infiltration scores of 23 immune cell types in TCGA. (G) Comparison of immune infiltration scores of 23 immune cell types between CHST2-high and CHST2-low groups in the TCGA cohort. (H) Comparison of type I interferon scores between CHST2-positive and CHST2-negative epithelial cells in TNBC samples at the single-cell level. * *p* <0.05; ** *p* <0.01; *** *p* <0.001; **** *p* <0.0001.

We next quantified immune and stromal components using the ESTIMATE framework. CHST2 expression correlated positively with immune score and overall ESTIMATE score across TCGA samples (Fig. 4C–E). When analyzed by subtype, this association with immune score was preserved in both TNBC and non-TNBC tumors. In contrast, correlation with stromal score was evident in non-TNBC tumors but minimal in TNBC, pointing to subtype-specific differences in the microenvironment linked to CHST2 expression.

Consistent with these findings, CHST2 expression positively correlated with multiple immune cell populations across adaptive and innate compartments (Fig. 4F). CHST2-high tumors displayed increased infiltration scores for several immune subsets compared with CHST2-low tumors (Fig. 4G).

Single-cell transcriptomic analysis indicated that CHST2 expression was largely confined to epithelial tumor cells, with minimal expression in immune or stromal populations. Within TNBC epithelial cells, CHST2-positive cells exhibited higher type I interferon response scores than CHST2-negative cells (Fig. 4H), linking CHST2 expression to interferon-related programs at the tumor cell level.

These associations were independently validated in the GSE96058 cohort, where CHST2-high tumors similarly showed enrichment of immune-related pathways, positive correlations with immune and ESTIMATE scores, and increased immune infiltration (Fig. S4A–G), supporting the reproducibility of the CHST2–immune association across cohorts.

### CHST2 enhances migratory and invasive behavior of TNBC cells without affecting proliferation

To define the functional contribution of CHST2 to TNBC cell behavior, CHST2 expression was manipulated in MDA-MB-231 cells by lentiviral-mediated knockdown or overexpression. Efficient CHST2 knockdown and overexpression were confirmed at both the mRNA and protein levels by RT–qPCR and Western blotting (Fig. 5A–B, D–E).

**Figure 5.**
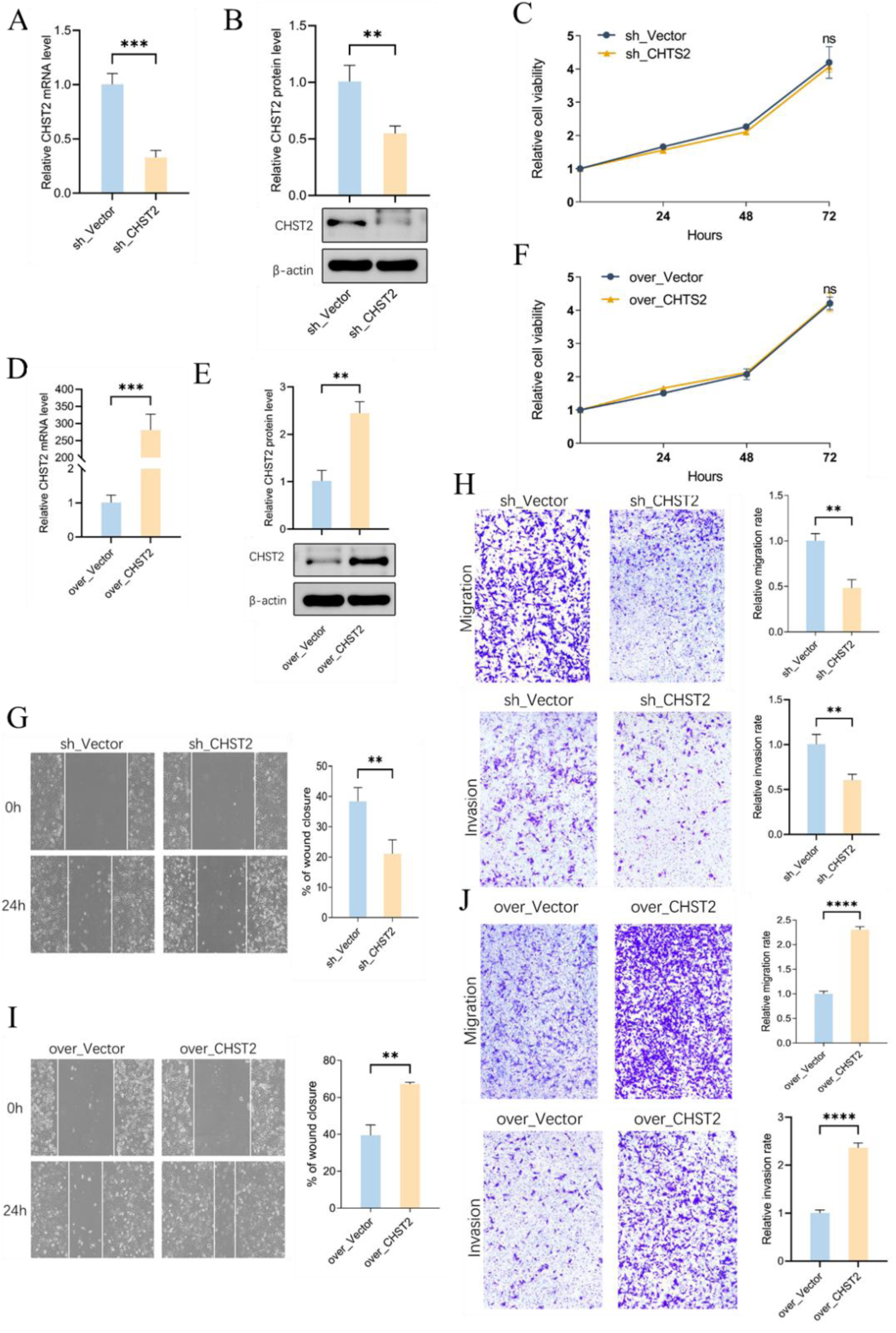
CHST2 promotes migration and invasion of TNBC cells. (A) RT–qPCR analysis of CHST2 mRNA expression in MDA-MB-231 cells transfected with control vector or CHST2 knockdown constructs. (B) Western blot analysis of CHST2 protein levels in control and CHST2-knockdown cells. (C) Cell proliferation assay in control and CHST2-knockdown cells. (D) RT–qPCR analysis of CHST2 mRNA expression cells transfected with control vector or CHST2 overexpression constructs. (E) Western blot analysis of CHST2 protein levels in control and CHST2-overexpressing cells. (F) Cell proliferation assay in control and CHST2-overexpressing cells. (G–H) Representative images of wound-healing assays (G) and Transwell migration assays (H) in control and CHST2-knockdown cells. (I–J) Representative images of wound-healing assays (I) and Transwell migration assays (J) in control and CHST2-overexpressing cells. Data are presented as mean ± SD from three independent experiments. ns, *p* ≥0.05; * *p* <0.05; ** *p* <0.01; *** *p* <0.001; **** *p* <0.0001.

Then, we firstly examined whether CHST2 influences cell proliferation. CCK-8 assays showed no significant differences in growth rates between control cells and cells with CHST2 knockdown or overexpression (Fig. 5C, F), indicating that CHST2 does not affect TNBC cell proliferation under these conditions. In contrast, CHST2 exerted a pronounced effect on cell motility. In wound-healing assays, CHST2 knockdown markedly delayed scratch closure, whereas CHST2 overexpression accelerated wound closure compared with control cells (Fig. 5G, I). Consistently, Transwell assays indicated that both migratory and invasive capacities were significantly reduced upon CHST2 depletion and enhanced following CHST2 overexpression (Fig. 5H, J).

Taken together, these results indicate that CHST2 selectively promotes migration and invasion of TNBC cells without affecting proliferative capacity, supporting a specific role for CHST2 in regulating tumor cell motility rather than growth.

### CHST2-driven migration and invasion partially require its catalytic activity

To determine whether the pro-migratory and pro-invasive effects of CHST2 depend on its enzymatic function, we generated a point mutant of CHST2 in which the conserved catalytic residue Asn475 was substituted with alanine (N475A) (Fig. 6A). This residue has been implicated in sulfotransferase activity in previous studies. Wild-type CHST2 and the N475A mutant were expressed in MDA-MB-231 cells in parallel. Immunoblotting confirmed comparable protein expression levels among empty vector, wild-type CHST2, and CHST2-N475A groups (Fig. 6B). Notably, CHST2-N475A exhibited a reduced apparent molecular weight, consistent with impaired post-translational modification, as previously reported.

**Figure 6.**
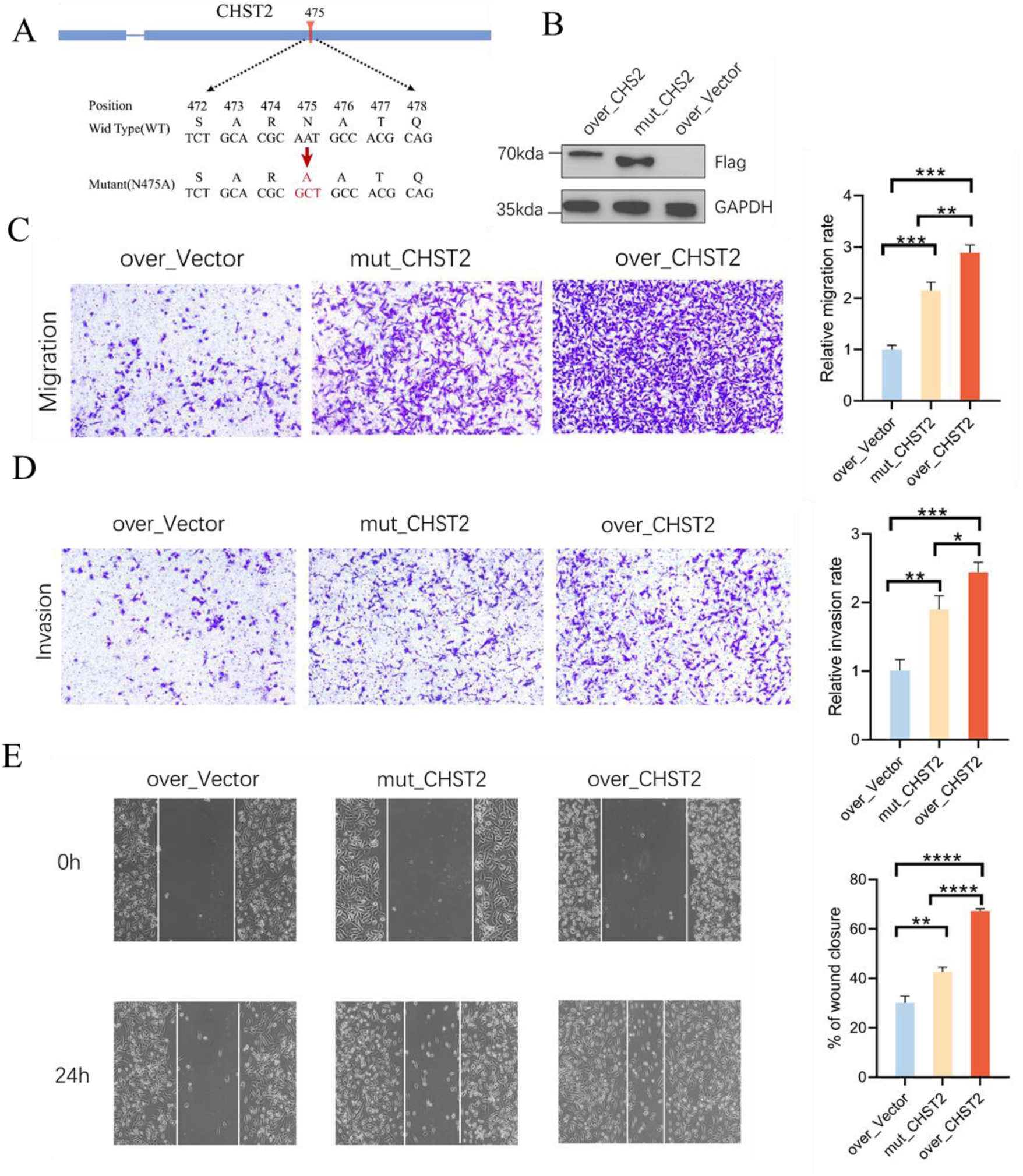
N475 is required for full CHST2-mediated migratory and invasive activity. (A) Schematic of the CHST2 N475A point mutation.****(B) Western blot analysis of CHST2 protein expression in cells transfected with empty vector, wild-type (WT) CHST2, or CHST2-N475A. (C–E) Representative images of Transwell migration (C), invasion (D), and wound-healing assays (E) in cells expressing empty vector, WT CHST2, or CHST2-N475A. Data are presented as mean ± SD from three independent experiments. ns, *p* ≥0.05; * *p* <0.05; ** *p* <0.01; *** *p* <0.001; **** *p* <0.0001.

We next assessed cell motility using Transwell-based assays. Consistent with the results shown in Fig. 5, wild-type CHST2 significantly enhanced both migration and invasion compared with vector controls, whereas the N475A mutant showed a markedly reduced effect (Fig. 6C–D). Quantitative analyses suggested a graded response, with maximal motility observed in wild-type CHST2–expressing cells, intermediate activity in N475A-expressing cells, and minimal activity in control cells.

A similar pattern was observed in wound-healing assays. Wild-type CHST2 substantially accelerated wound closure, whereas the N475A mutation significantly blunted this effect without fully abolishing it (Fig. 6E). Together, these data indicate that an intact catalytic residue is required for the full pro-migratory and pro-invasive activity of CHST2, while residual effects persist in the absence of catalytic integrity.

Taken together, the gain- and loss-of-function analyses suggest that CHST2 enhances TNBC cell migration and invasion independently of proliferation, and that this pro-motile activity is partially dependent on its catalytic function, supporting a model in which CHST2 promotes aggressive tumor cell behavior through both activity-dependent and activity-independent mechanisms.

### Promoter hypomethylation underlies CHST2 activation in TNBC

To define the epigenetic basis of subtype-specific CHST2 expression, we examined DNA methylation across its promoter region. CpG island analysis identified a CpG-rich promoter segment (Fig. 7A). TCGA methylation data showed significantly lower methylation levels at multiple CpG sites within the CHST2 promoter in TNBC compared with non-TNBC tumors (Fig. 7B). This pattern was recapitulated in breast cancer cell lines. Genome-wide methylation profiling revealed predominant promoter hypomethylation in TNBC cell lines, whereas non-TNBC lines exhibited marked hypermethylation at the same locus (Fig. 7D).

**Figure 7.**
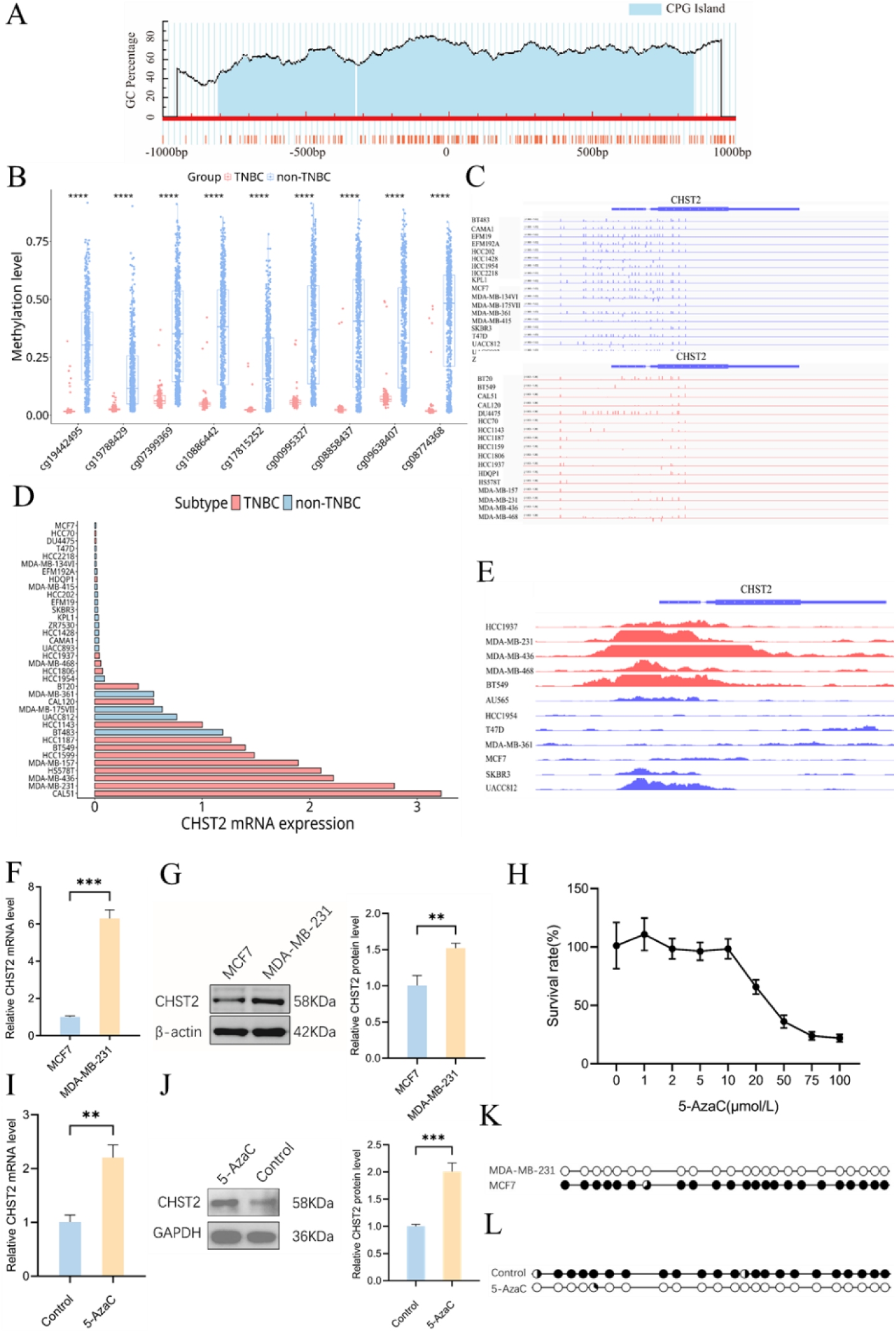
Epigenetic regulation of CHST2 in TNBC and non-TNBC cells. (A) Schematic representation of CpG distribution in the CHST2 promoter region. (B) Comparison of CHST2 promoter CpG methylation between TNBC and non-TNBC samples from the TCGA. (C) CpG methylation status of the CHST2 promoter in TNBC and non-TNBC cell lines. Red indicates TNBC cell lines; blue indicates non-TNBC cell lines. (D) CHST2 expression levels across different breast cancer cell lines. (E) Distribution of H3K27ac histone modification signals at the CHST2 promoter across breast cancer cell lines. Color coding as in (C). RT-qPCR (F) and Western blot (G) analysis of CHST2 expression in MDA-MB-231 and MCF7 cells. (H) Bisulfite sequencing PCR analysis of CHST2 promoter methylation in MDA-MB-231 and MCF7 cells. RT-qPCR (I) and Western blot (J) analysis of CHST2 expression in MCF7 cells following 5-AzaC treatment. (K) Effects of different concentrations of 5-AzaC on MCF7 cell viability. (L) BSP analysis of CHST2 promoter CpG island methylation in MCF7 control and 5-AzaC–treated cells. Each circle represents a CpG site; black indicates methylated, white indicates unmethylated. Data are presented as mean ± SD from three independent experiments. Statistical significance: ns, *p* ≥0.05; * *p* <0.05; ** *p* <0.01; *** *p* <0.001; **** *p* <0.0001.

We next assessed chromatin activation status. H3K27ac ChIP–seq data demonstrated strong enrichment of this active histone mark at the CHST2 promoter in TNBC cell lines, with substantially weaker signals in non-TNBC lines (Fig. 7C), consistent with an open chromatin configuration in TNBC.

To directly link promoter methylation with CHST2 expression, we compared MDA-MB-231 (TNBC) and MCF7 (non-TNBC) cells. CHST2 expression was markedly higher in MDA-MB-231 cells at both mRNA and protein levels (Fig. 7F–G). Bisulfite sequencing confirmed extensive promoter hypomethylation in MDA-MB-231 cells and dense CpG methylation in MCF7 cells (Fig. 7H).

Pharmacological demethylation further substantiated this relationship. Treatment of MCF7 cells with the DNA methyltransferase inhibitor 5-aza-2′-deoxycytidine resulted in a marked reduction of methylation levels at the CHST2 promoter (Fig. 7L) and led to a pronounced increase in CHST2 mRNA and protein expression (Fig. 7I–J), without affecting viability under the tested conditions (Fig. 7H), indicating that promoter methylation directly constrains CHST2 transcription.

Collectively, these data identify promoter hypomethylation as a key epigenetic determinant of CHST2 activation in TNBC.

## Discussion

While widespread epigenetic alterations have been described in triple-negative breast cancer, direct links between specific DNA methylation events and functionally relevant oncogenic programs remain limited(25, 26). In this study, through integrated epigenomic, transcriptomic, and functional analyses, we identify carbohydrate sulfotransferase 2 (CHST2) as an epigenetically activated regulator in TNBC and suggest that promoter hypomethylation is a key determinant of its aberrant expression and pro-invasive function.

In both primary TNBC tumors and breast cancer cell line models, genes exhibiting promoter hypomethylation accompanied by increased expression constituted the most prevalent methylation–expression category(27). Although the specific gene sets showed limited overlap between systems, this pattern-level concordance suggests that loss of promoter methylation represents a recurrent mechanism contributing to transcriptional reprogramming in TNBC(27). Within this landscape, CHST2 consistently displayed a strong inverse relationship between promoter methylation and transcript abundance. Notably, among members of the carbohydrate sulfotransferase family, CHST2 exhibited the most robust and reproducible epigenetic activation across independent datasets, arguing against a nonspecific consequence of global DNA hypomethylation. Instead, these findings suggest that CHST2 activation reflects selective epigenetic deregulation with functional relevance in TNBC progression.

Single-cell transcriptomic analyses further revealed marked cell type– and subtype-specific patterns of CHST2 expression. In TNBC, CHST2 was predominantly enriched in malignant epithelial cells, whereas in non-TNBC tumors its expression was largely confined to endothelial compartments. This reciprocal distribution suggests context-dependent biological roles for CHST2 across breast cancer subtypes(28, 29). In TNBC, epithelial-specific upregulation likely represents a tumor cell–intrinsic adaptation associated with enhanced migratory capacity and invasive potential(30, 31). In contrast, CHST2 expression in endothelial and stromal cells of non-TNBC tumors is consistent with its established involvement in vascular function and immune cell trafficking, highlighting distinct functional roles across breast cancer subtypes.

Consistent with this notion, functional assays indicated that CHST2 selectively promotes migration and invasion of TNBC cells without significantly affecting proliferation. Such dissociation between proliferative capacity and invasive behavior has been reported in aggressive breast cancers and is thought to reflect the biological separation of metastatic competence from tumor growth(12). In this context, CHST2 appears to contribute primarily to invasive plasticity, in line with its positive association with mesenchymal and EMT-related transcriptional programs observed in TNBC tumors.

Mechanistically, the pro-migratory effects of CHST2 were largely dependent on its sulfotransferase activity(12, 32, 33). Disruption of the catalytic domain markedly attenuated CHST2-driven migration and invasion, supporting a role for altered glycan sulfation in promoting invasive behavior. The incomplete abrogation of motility observed with the catalytic mutant suggests that CHST2 may also exert activity-independent effects, potentially through interactions with other components of the glycosylation machinery. Identification of specific sulfated substrates and downstream signaling pathways will be necessary to fully elucidate how CHST2-mediated glycan remodeling contributes to TNBC progression.

Beyond tumor cell–intrinsic effects, elevated CHST2 expression was associated with broad immune-related transcriptional changes and increased immune infiltration in TNBC tumors. Although CHST2 expression itself was largely confined to malignant epithelial cells, CHST2-high tumors displayed immune features indicative of both immune activation and immunosuppressive signaling. Such immune-enriched yet clinically aggressive phenotypes have been reported in TNBC and underscore the complex and context-dependent nature of tumor–immune interactions in this disease. Recent studies have highlighted that tumor-associated glycan remodeling can actively shape immune responses within the tumor microenvironment (34). Importantly, these observations suggest an association rather than a direct regulatory role of CHST2 in immune modulation. Given the established involvement of glycan sulfation in immune receptor engagement, leukocyte trafficking, and cytokine signaling(35), it is plausible that CHST2-driven alterations in the tumor glycome may indirectly contribute to immune remodeling within the tumor microenvironment(36). However, the precise mechanisms linking CHST2-mediated sulfation to immune regulation remain to be elucidated.

At the epigenetic level, our findings provide direct evidence that promoter hypomethylation is a primary mechanism driving CHST2 activation in TNBC. Bisulfite sequencing, pharmacological demethylation, and transcriptional analyses consistently suggested an inverse relationship between promoter methylation and CHST2 expression, supporting a causal link between DNA methylation deregulation and activation of a glycan-modifying enzyme that promotes invasive behavior. These results reinforce the concept that epigenetic alterations can directly engage noncanonical oncogenic pathways in aggressive cancers(37).

From a clinical perspective, elevated CHST2 expression has been linked to unfavorable patient outcomes in independent clinical cohorts, suggesting a potential association with aggressive disease behavior. While this relationship was not uniformly observed across all publicly available datasets(38), these findings raise the possibility that CHST2 expression or promoter methylation status may carry prognostic relevance in specific clinical contexts. Further validation in well-annotated TNBC cohorts will be required to clarify its clinical utility.

Taken together, this study identifies CHST2 as an epigenetically activated regulator of invasive behavior in TNBC and links promoter hypomethylation to sulfotransferase-dependent tumor cell motility. Our findings highlight an underappreciated connection between epigenetic deregulation and glycan modification in TNBC and provide a rationale for further exploration of epigenetically driven sulfation pathways as potential therapeutic targets in aggressive breast cancer.

## Acknowledgements

This work was supported by the Fundamental Research Funds for the Central Universities and Hubei Key Laboratory of Genetic Regulation and Integrative Biology.

## Authorship contribution statement

Shukun Qu: Data curation, Visualization, Validation, Software, Methodology, Investigation, Conceptualization; Kangning Li: Validation, Investigation. Lili Fan: Investigation, Visualization. Chenxue Miao: Methodology; Weiyu Wang: Writing-review & editing; Xu Wang: Methodology; Guang Song: Conceptualization, Supervision, Original draft, Formal analysis, Writing-original draft, review & editing, Funding acquisition.

## Data availability statement

The data that support the findings of this study are available from the corresponding author upon reasonable request.

