## Supplementary figures for "Epigenetic activation of CHST2 by promoter hypomethylation promotes progression of triple-negative breast cancer"

### Supplementary Fig. 1

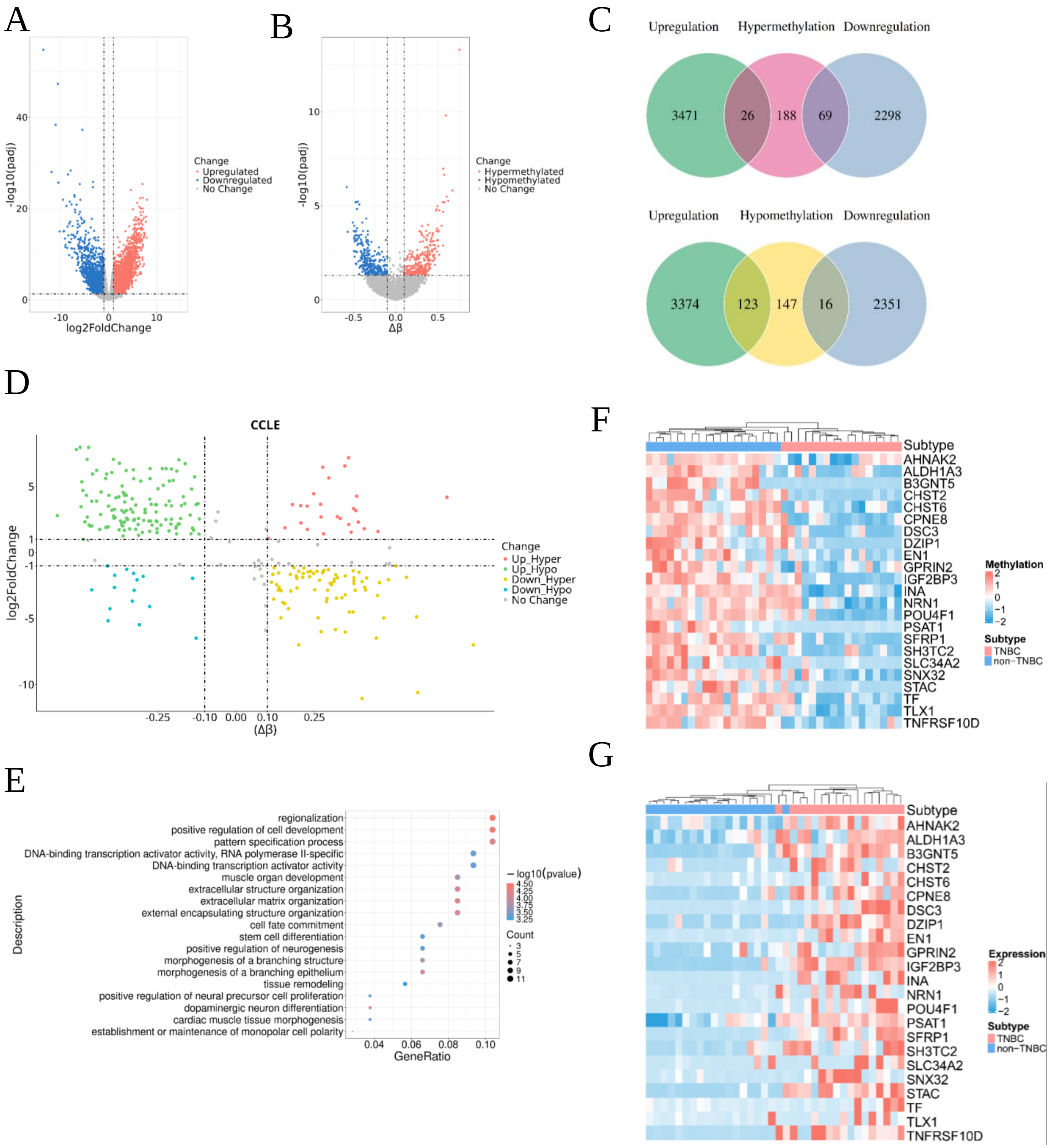


**Supplementary Fig. 1 Promoter hypomethylation–associated gene activation in TNBC cell lines in CCLE**

(A) Volcano plot of differentially expressed genes (DEGs) between TNBC and non-TNBC cell lines in the CCLE dataset. Red and blue dots indicate significantly upregulated and downregulated genes in TNBC. (B) Volcano plot of gene-level promoter methylation differences between TNBC and non-TNBC cell lines in CCLE. Red dots denote hypermethylated promoters and blue dots denote hypomethylated promoters in TNBC. (C) Venn diagrams showing the overlap between genes with altered promoter methylation and upregulated or downregulated genes in the TCGA cohort. (D) Integrated analysis of DEGs and genes with differential promoter methylation (DMGs) in CCLE, classifying genes into five regulatory patterns: Up_Hyper, Up_Hypo, Down_Hyper, Down_Hypo, and No Change. (E) Gene Ontology (GO) enrichment analysis of genes exhibiting the Up_Hypo pattern in CCLE. (F–G) Heatmaps showing expression levels (F) and promoter methylation levels (G) of 23 key Up_Hypo genes in TNBC and non-TNBC cell lines from the CCLE dataset.

### Supplementary Fig. 2

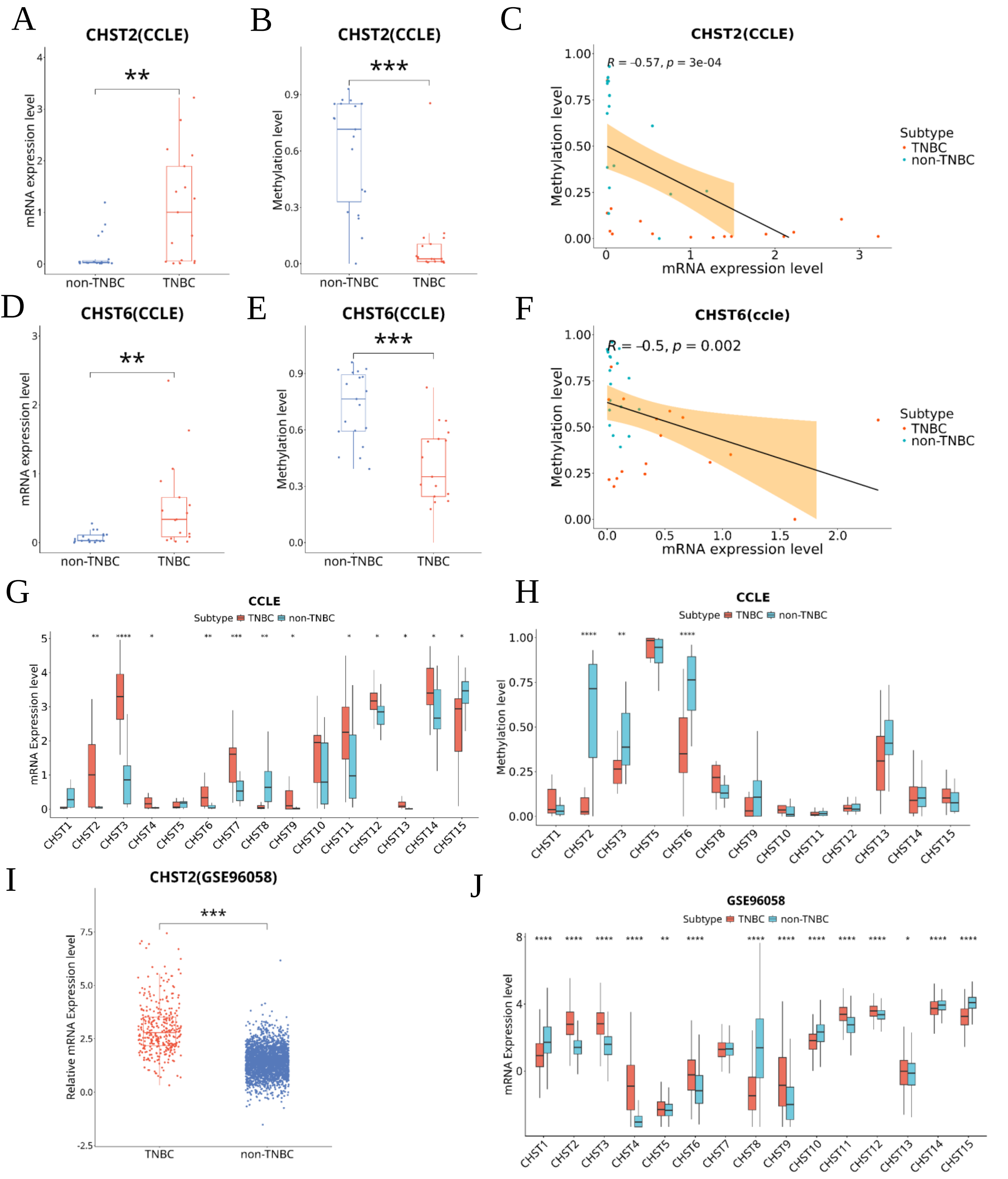


**Supplementary Fig. 2 Epigenetic features and clinical relevance of CHST family members in CCLE and GSE96058 datasets.** (A–B) Box plots showing CHST2 mRNA expression (A) and its promoter methylation (B) in TNBC and non-TNBC cell lines from the CCLE dataset. (C) Correlation analysis between CHST2 mRNA expression and its promoter methylation in CCLE cell lines. (D–E) Box plots showing CHST6 mRNA expression (D) and its promoter methylation (E) in TNBC and non-TNBC cell lines from CCLE. (F) Correlation analysis between CHST6 mRNA expression and its promoter methylation in CCLE cell lines. (G) Box plots summarizing differential expression of CHST family members between TNBC and non-TNBC cell lines in CCLE. (H) Box plots showing promoter methylation differences of CHST family genes between TNBC and non-TNBC cell lines in CCLE. (I) CHST2 expression levels in TNBC and non-TNBC breast cancer samples from the GSE96058 cohort. (J) Differential expression of CHST family genes between TNBC and non-TNBC samples in the GSE96058 cohort.

NS $P \geq 0.05$; * $p < 0.05$; ** $p < 0.01$; *** $p < 0.001$; **** $p < 0.0001$.

### Supplementary Fig. 3

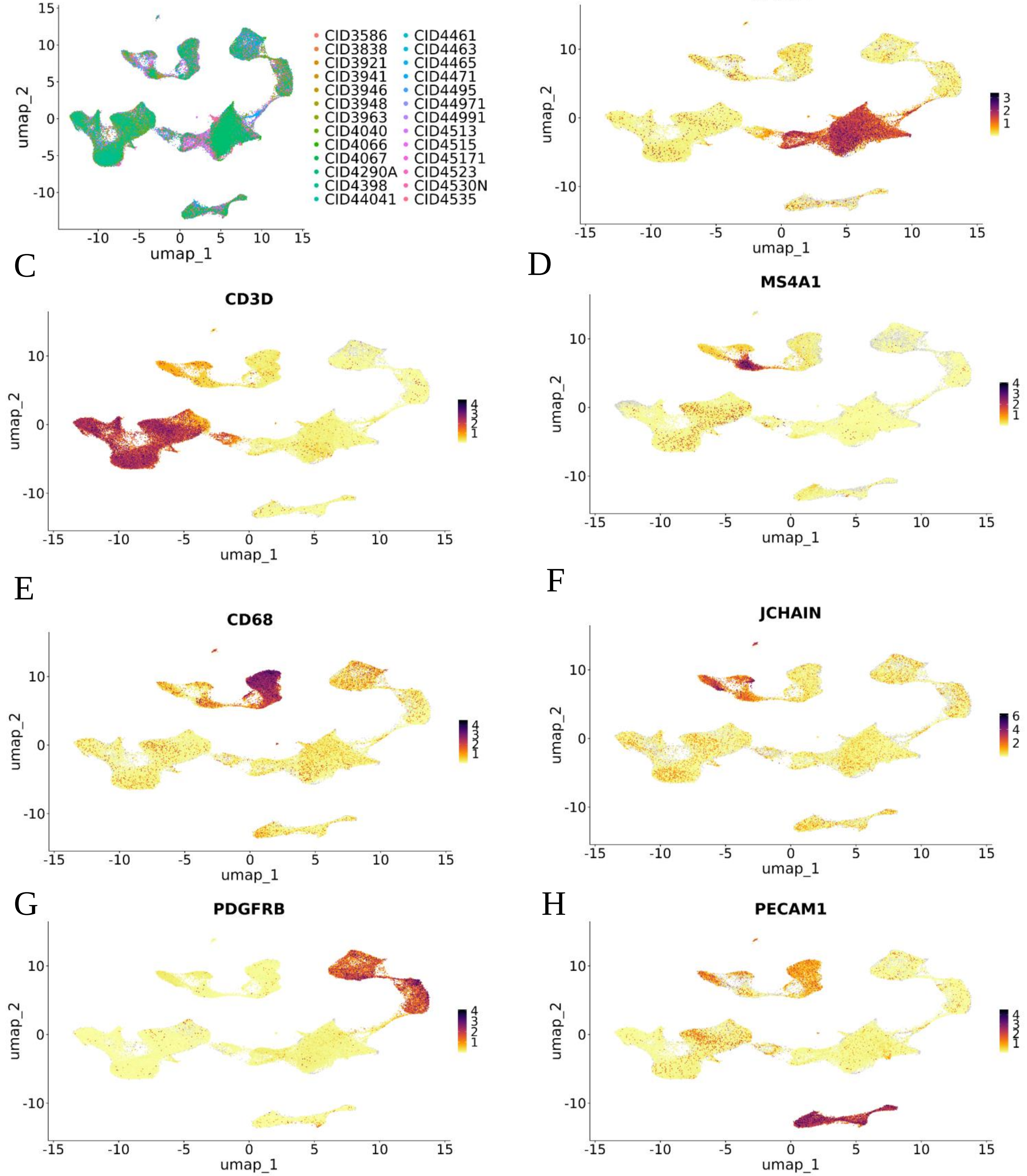


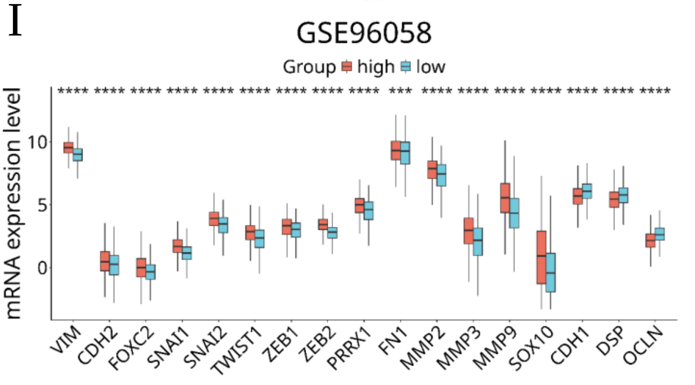


**Figure S3. Batch correction, cell-type annotation, and validation of CHST2-associated EMT signatures.**

(A) Batch-corrected distribution of breast cancer samples. (B–H) Expression of canonical marker genes used for cell-type annotation. (I) Differential expression of EMT-related genes between CHST2-high and CHST2-low groups in the GSE96058 dataset.

**Supplementary Fig. 4**

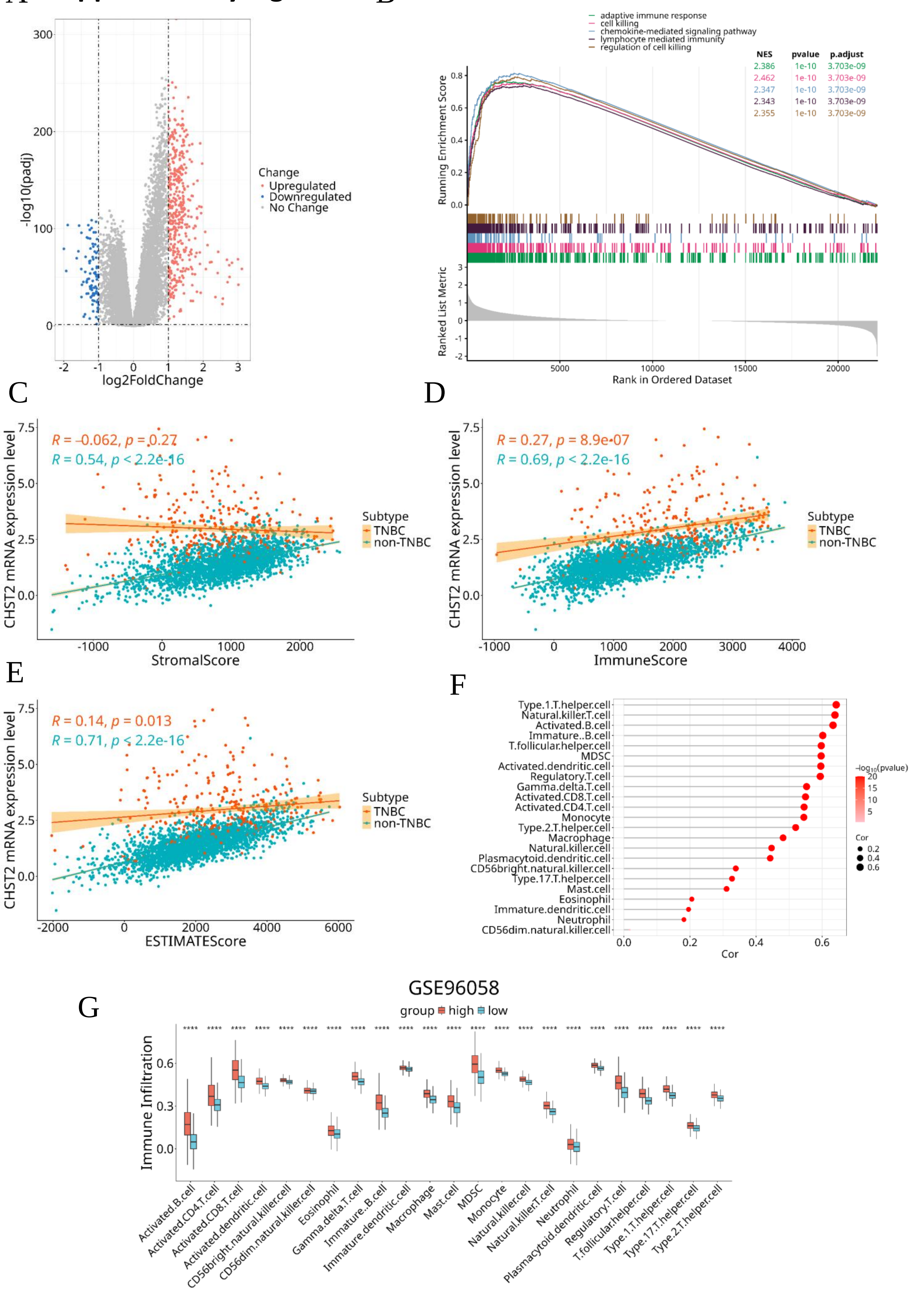

**Fig. S4 Association of CHST2 expression with tumor microenvironment characteristics and immune infiltration in the GSE96058 cohort.**

(A) Differentially expressed genes between CHST2-high and CHST2-low groups in the GSE96058 cohort. Samples were stratified according to the median CHST2 expression level. (B) Gene set enrichment analysis of differentially expressed genes between CHST2-high and CHST2-low groups in GSE96058, showing the top five pathways ranked by normalized enrichment score (C–E) Correlation analysis between CHST2 expression and stromal score (C), immune score (D), and ESTIMATE score (E) in GSE96058 samples. Each dot represents one tumor sample; TNBC and non-TNBC samples are indicated by different colors. Correlation coefficients and *P* values were calculated using Spearman's correlation analysis. (F) Lollipop plot showing correlations between CHST2 expression and infiltration scores of 23 immune cell types in the GSE96058 cohort. (G) Comparison of immune infiltration scores of 23 immune cell types between CHST2-high and CHST2-low groups in the GSE96058 cohort.
